# Local design principles at hippocampal synapses revealed by an energy-information trade-off

**DOI:** 10.1101/748400

**Authors:** Gaurang Mahajan, Suhita Nadkarni

## Abstract

Synapses across different brain regions display distinct structure-function relationships. We investigate the interplay of fundamental design principles that shape the transmission properties of the excitatory CA3-CA1 pyramidal cell connection, a prototypic synapse for studying the mechanisms of learning in the hippocampus. This small synapse is characterized by probabilistic release of transmitter, which is markedly facilitated in response to naturally occurring trains of action potentials. Based on a physiologically realistic computational model of the CA3 presynaptic terminal, we show how unreliability and short-term dynamics of vesicle release work together to regulate the trade-off of information transfer versus energy use. We propose that individual CA3-CA1 synapses are designed to operate at close to maximum possible capacity of information transfer in an efficient manner. Experimental measurements reveal a wide range of vesicle release probabilities at hippocampal synapses, which may be a necessary consequence of long-term plasticity and homeostatic mechanisms that manifest as presynaptic modifications of release probability. We show that the timescales and magnitude of short-term plasticity render synaptic information transfer nearly independent of differences in release probability. Thus, individual synapses transmit optimally while maintaining a heterogeneous distribution of presynaptic strengths indicative of synaptically-encoded memory representations. Our results support the view that organizing principles that are evident on higher scales of neural organization percolate down to the design of an individual synapse.

## 1. INTRODUCTION

Signal transmission at chemical synapses accounts for a significant proportion of metabolic costs during normal neural activity in the mammalian brain^1^. Understanding the role of competing demands imposed by energy consumption and information processing in shaping nervous systems has been an enduring question in neuroscience research^2–5^; one may then ask if the notion of energetic efficiency trickles down to the level of individual synapses. In this context, failures of transmitter release, while unsuccessful in relaying presynaptic action potentials, may help conserve synaptic resources by lowering average release rates. Indeed, probabilistic release is a characterizing feature found across a number of synapses^6^, and a fundamental source of stochasticity in neural dynamics^7^. Previous studies have suggested that synaptic failures support both efficient neuronal coding^8^ and communication between neurons^9^, but these studies did not include the effect of use-dependent short-term plasticity (STP) that typically accompanies probabilistic release and can significantly modulate the time course of synaptic responses to natural activity patterns^10–12^.

Excitatory CA3-CA1 pyramidal cell connections, a crucial component of the hippocampal circuitry engaged during spatial navigation and implicated in important forms of learning^13–16^, provide a distinctive example of low release probability synapses^17^. Low transmission rates for single spikes are contrasted with strong enhancement of release probabilities in response to natural stimuli^18^; this short-term facilitation (STF) occurs over timescales of milliseconds to seconds. Dynamic CA3-CA1 synapses were proposed to be optimally designed for conveying information on spike times in short bursts occurring at physiologically relevant frequencies^19^. However, the concomitant energy costs associated with vesicle release and recycling supporting this form of transmission are not known.

Here, we use a computational model to investigate the relevance of energetic constraints to design and function of hippocampal synapses that are characterized by low initial release probabilities but marked activity-dependent STP. Previous studies of information transmission at cortical synapses considered short-time dynamics arising from vesicle depletion alone^20–23^, or made simplifying model assumptions about presynaptic organization (e.g., availability of at most one vesicle per release site)^9,24^, limiting their physiological relevance for describing facilitating hippocampal synapses. Another distinguishing feature of our study is that we do not ascribe a notion of information to ‘a’ spike as is often done^9,21^, as it is unclear if every presynaptic spike can be assigned meaning at the CA3-CA1 synapse. In the hippocampus, neural information may be encoded in changing firing rates rather than the precise timing of individual spikes. A relevant example is provided by the selective activation of specific subsets of pyramidal cells whenever the animal enters their preferred spatial location^25^, and such brief increases in firing punctuating a low-activity background are regarded as units of information^26^. This suggests that, instead of a ‘spike-centric’ approach assessing how reliably synapses with dynamic strengths convey presynaptic spike times to the target cell, it may be more meaningful to work with a notion of information that directly relates to a physiologically identifiable temporal ‘signal’ encoded in the irregular firing activity of the presynaptic neuron.

Vesicle release properties are seen to vary widely across synapses, being tuned to the functional demands of the circuits in which they are embedded^27–29^. Thus, addressing design principles on a general level is impeded by the diversity of synapse types found in the nervous system and the neural activity patterns that they process. How a small hippocampal synapse defined by low release probability and a limited pool of available vesicles, equipped with short-term plasticity, regulates the local balance between reliability and economy of signaling in a physiological setting has not been addressed thus far. Our model includes relevant biological details and characterizes the role of activity-dependent, short-term release dynamics in modulating the transmission of rate-coded presynaptic signals. CA3 synaptic populations display considerable heterogeneity in their transmitter release properties^30–32^, and we particularly sought to address how these differences among synapses impact their ability to relay information-carrying spike trains. We show that the CA3-CA1 synapse operates in a regime that maintains low energy costs while maximizing information dynamically via STP for the entire heterogeneous population of intrinsic release probabilities seen at these synapses.

## 2. METHODS

Experimental methods have provided valuable information about the ultratructural organization and distribution of dynamic properties at CA3-CA1 presynaptic terminals^30,33^. Individual CA3 synaptic boutons typically have a single active zone^33^, where glutamate release occurs in a probabilistic manner^17^ and shows a complex mix of use-dependent depression due to refractoriness in vesicle recovery and rapid calcium-mediated facilitation of the release machinery^29,34^. We adopted a mathematical description of this synapse which captures key attributes of its short-time release dynamics, and quantified through numerical simulations its responses to irregular spike trains mimicking naturally occurring presynaptic cell activity. The model details and setup for our analysis are briefly described below.

### Estimating synaptic information rates and efficiency

Experimental recordings from rodent hippocampus suggest that individual pyramidal cells in area CA3 show location-specific (place cell) firing during free exploration^35^. Further it has been argued that variability seen in these brief increases in firing during individual passes through the preferred location may encode additional attributes such as aspects of the animal’s trajectory^36,37^, its motion relative to goal direction^38^, variable attentional state of the animal^39,40^, modulation by contextual cues^41,42^ (besides simply the variable duration, or equivalently, the running speed, at each pass), etc. These observations motivate our model wherein we consider synaptic processing of integrated spatial and contextual signals represented by the timing and size (number of spikes) of place field discharges of the CA3 neuron. We thus define an effective ‘input’ signal, *ϕ*(t), underlying the variable firing activity of the presynaptic cell which is assumed to be zero everywhere except whenever the trajectory crosses the cells preferred location. Every pass through the place field is associated with a non-zero value of *ϕ, ϕ*_*i*_, which in our model is sampled from a uniform distribution on the interval [*ϕ*_min_, *ϕ*_max_]. This range is adjusted to be compatible with the statistics of experimentally recorded hippocampal spike train data (described below).

Individual place field passes are assumed to be uncorrelated in time and occur sparsely, at a mean rate of *r*_*s*_ s^−1^. In order to estimate information rate carried by *ϕ*(t), time is uniformly divided into sufficiently short steps of Δt (≪ 1*/r*_*s*_), which is taken to be the (fixed) duration of every pass; every time step is thus treated as an independent realization of the probability distribution of *ϕ*, P(*ϕ*), that is determined by the value of *r*_*s*_. The time-averaged, discretized, entropy rate of the input signal is quantified in the usual manner using Shannon’s measure^43^:

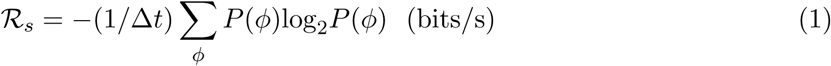

where the sum runs over all values that *ϕ* can assume, P(*ϕ*) here being approximated by a discrete distribution over n_*ϕ*_ possible states (set to 20 in our analysis). The value of *ϕ* during every place field pass determines the corresponding burst size; the number of spikes comprising every burst is thus given by a second, conditional, Poisson distribution with mean *λ*(*ϕ*) (the exact form of which depends on the specific interpretation of the *ϕ* variable; see below), and these spikes are assumed to occur at random times within the corresponding pass of duration Δt. To be consistent with experimental data, a small amount of ‘noise’ is also added to the system, modeled as a constant background presynaptic spiking rate of *r*_*n*_ *s*^−1^ (this spiking is uncorrelated with the spatial context and could arise, say, from synaptic or channel noise).

The synaptic response to presynaptic spike patterns consists of a sequence of evoked transmitter release events, and we quantify how well this discrete temporal sequence conveys the temporal modulation of the signal *ϕ* underlying the irregular firing behavior of the presynaptic neuron. In analogy with the input, we approximate the CA3 synaptic output by binning releases occurring within every time step Δt. Thus, every burst evokes a variable number of release events, *n*_*r*_, and the coarse-grained response profile is given by a sequence of *n*_*r*_ values (one number per Δt step, and that are assumed to produce graded postsynaptic responses via temporal summation of EPSPs or NMDA receptor-gated Ca^2+^ transients). Under the assumption of low noise level, synaptic transmission is characterized by the stationary joint probability distribution P(*ϕ, n*_*r*_) ≡ P(*ϕ*)P(*n*_*r*_|*ϕ*) which is (implicitly) sampled at every time step in our simulations. The conditional distribution P(*n*_*r*_|*ϕ*) is governed by the form of the synaptic dynamics used in the model and encapsulates the effects of STP. Synaptic responses to successive bursts can be considered to be uncorrelated, which is a valid approximation when the typical interval between place field passes is longer than the slowest timescale in the model of synaptic dynamics (this is set by the recovery rate of the release-ready vesicle pool; see below). We characterize the fidelity of information transmission at an individual synapse in terms of a discretized version of the average mutual information rate (*R*_mutual_), a standard nonparametric measure of statistical relatedness of two variables, which can be expressed as a difference between the total entropy of the synaptic response and the *noise* entropy^43^:

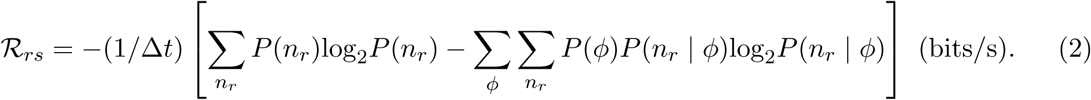

This rate is numerically estimated from the pooled data from the simulation run for sufficiently long duration.

Following earlier studies^9,21^, local efficiency of the synaptic code is quantified in terms of the average number of releases per bit transmitted per synapse, and we use the measure *E*(*s*^−1^) = *N*_*r*_*/*(*ℛ*_*rs*_*/ℛ*_*s*_), where *N*_*r*_ denotes the mean rate of fusion events (averaged over every simulation run). *N*_*r*_ accounts for use of synaptic resources during signal transmission and also provides a proxy for the energetic costs of generating postsynaptic responses (membrane potential transients).

### Dynamical model of probabilistic synapses

Every synaptic release site is characterized by its basal spike-evoked transmission probability, 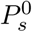, and maximum size of the docked pool of release-ready vesicles (RRP), *N*_max_. The synaptic release probability (*P*_*s*_) is distinct from the fusion probability *per docked vesicle* (*p*_*v*_), and under the assumption that docked vesicles can fuse independently of each other, the two are related by 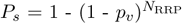, *N*_RRP_ being the instantaneous RRP size at the release site.

We model the synchronous component of vesicle release evoked by presynaptic spikes^30^, and assume that every spike can trigger release of at most one vesicle per synapse. This assumption of uniquantal release at glutamatergic CA3-CA1 synapses is compatible with experimental findings^34^ suggesting a refractory phase associated with fusion of a vesicle, which may inhibit subsequent release events in a short time period (∼10 ms) following the initial spike when the local calcium concentration at the release site is high. Following its release, every vesicle is assumed to be recovered independently, and this refilling is also modeled as a stochastic process with mean recovery timescale of *τ*_*r*_ per vesicle. During bouts of intense spiking activity, rapid depletion of the docked vesicle pool can occur, mediating a form of transient depression whose strength and duration are controlled by *τ*_*r*_ together with the resting pool size at the synapse. We set *τ*_*r*_ to 2 s, consistent with experimentally measured refilling rates at hippocampal synapses^44^.

Transient depression due to slow recycling of released vesicles is complemented by activity-dependent changes in vesicle fusion at the release site. We adopt a reduced kinetic model (Fig. 1A) to describe this short-term facilitation (STF) regulated by spike-driven calcium dynamics at the active zone, which follows from a number of previous studies of presynaptic plasticity^11,45^. The spike-triggered per-vesicle release probability, *p*_*v*_, is treated as a dynamical variable whose dependence on the presynaptic spiking history is governed by the following equation:

**FIG. 1:**
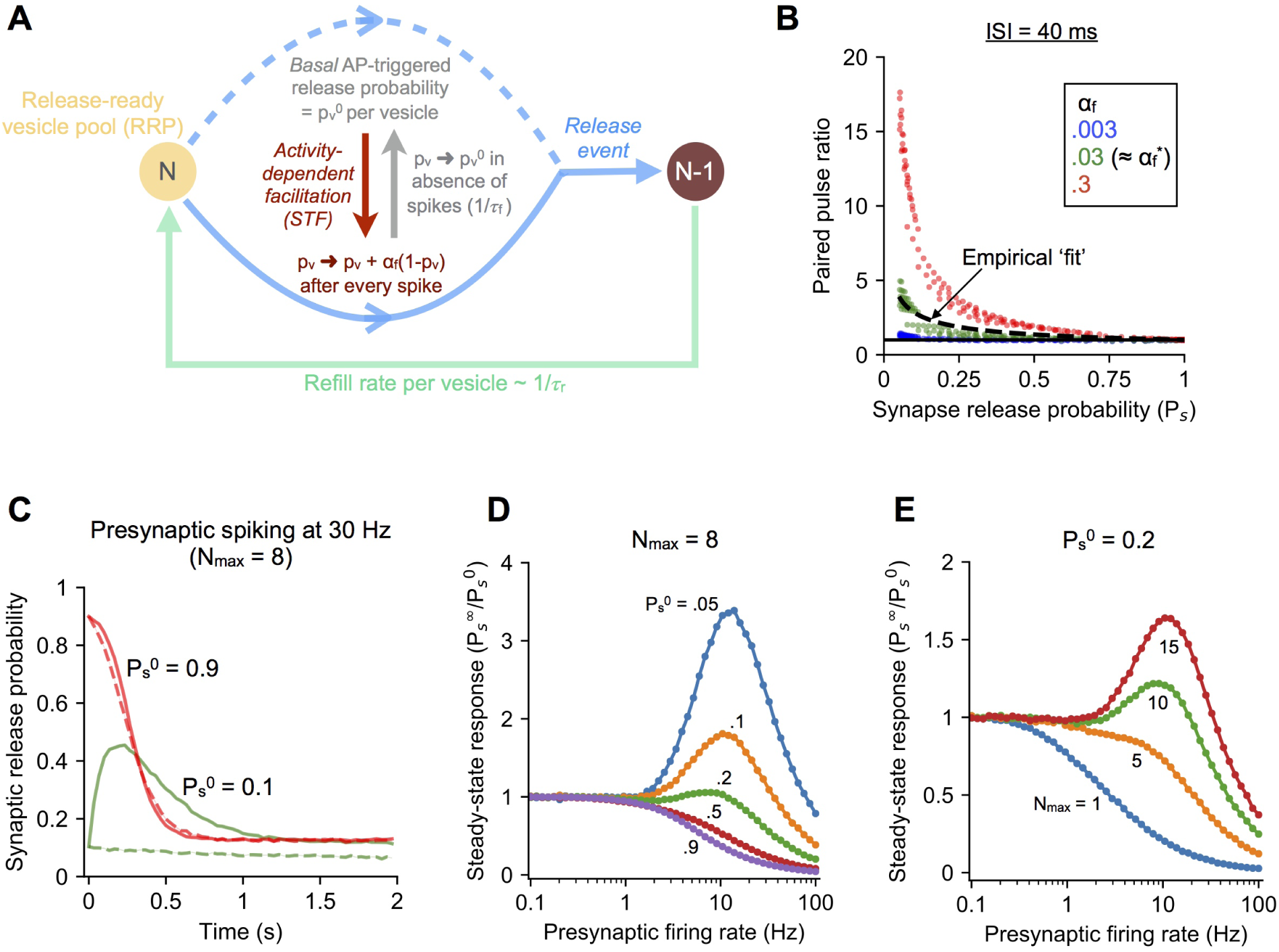
Modeling short-term plasticity (STP) at stochastic hippocampal synapses. **(A)** Outline of the reduced model of presynaptic STP used in the study, which includes activity-dependent facilitation of neurotransmitter release (STF) and depression due to slower recovery of released vesicles. STF is controlled by the dimensionless gain parameter *α*_*f*_. **(B)** Distribution of paired-pulse facilitation ratios (PPR) over a realistic range of RRP sizes (1-15) and basal per-vesicle release probabilities (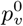 ranging from 10^−4^ to 1) in the STP model. Different levels of facilitation (*α*_*f*_) are represented by different colors. The dashed curve captures the empirical distribution of PPR values (from Dobrunz & Stevens, 1997), and the solid black line corresponds to PPR = 1. **(C)** Example of STP dynamics at realistic synapses with high 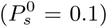 and low 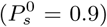 initial failure rates during response to regular presynaptic spiking at 30 Hz. Solid and dashed curves correspond to synapses with STF and lacking STF (constant *p*_*v*_), respectively. Results shown as mean *±* SEM over 10^4^ independent trials; *N*max= 8. **(D)** Regimes of facilitation and depression illustrated by the frequency dependence of normalized asymptotic/steady-state response for synapses with different basal failure rates (different colors). All synapses have the same maximum RRP size of 8. Results shown as mean *±* SEM over 10^4^ trials per parameter combination. **(E)** Dependence of synaptic filtering on the number of available vesicles illustrated by the frequency-response curves for synapses with the same initial release probability 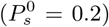 and varying maximum RRP size (different colors represent different *N*_max_). Results shown as mean *±* SEM over 10^4^ trials per parameter combination.

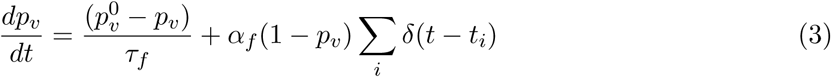

with the sum running over the set of all spike times. Thus, the arrival of every presynaptic spike increments the value of *p*_*v*_ by an amount proportional to the synaptic gain parameter *α*_*f*_, and the factor (1 - *p*_*v*_) ensures that *p*_*v*_, being a probability, does not exceed 1. The first term on the RHS of Eq. 1 describes the exponential relaxation of *p*_*v*_ to its baseline value 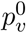 in the absence of spiking activity. Interspike intervals shorter than the facilitation time constant, *τ*_*f*_, are expected to induce strong enhancement of vesicle release. Following previous studies, we model a rapid form of STF with *τ*_*f*_ = 150 ms^44,46^. Longer-lasting forms of presynaptic plasticity such as augmentation are not considered here; these components are normally induced in experimental settings with sustained high frequency synaptic stimulation^10^, and unlikely to be of significance during the sparse, sporadic spiking activity observed in the physiological conditions modeled here.

The form of STF given by Eq. 3 implies a general trend of decreasing facilitation with increase in the basal synaptic release efficacy, which is supported by experimental recordings of individual hippocampal synapses^30,31^. In order to select a physiologically relevant value for the gain parameter *α*_*f*_, we refer to previous experimental data on paired-pulse stimulation at rat CA3-CA1 synapses^30^. This synaptic population displayed a broad range of baseline *P*_*s*_ values (∼0.05-1), and the release probabilities recorded in response to two spikes separated by a short interval (40 ms) yielded a distribution of paired pulse facilitation ratios (PPR) whose dependence on the initial synaptic release probability (*P*_s_) was well-fit by the relation 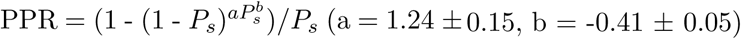. Using our simplified description of presynaptic dynamics, we analytically estimated the PPR in our model over a realistic range of basal per-vesicle release probabilities 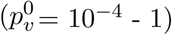 and RRP sizes (*N*_max_ = 1-15), and found the value of *α*_*f*_ for which this distribution of values was best fit by the above empirical model. The minimum mean-squared error was obtained at *α*_*f*_ *≈* 0.03. We thus set 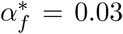 as the biological reference value of the gain parameter in our simulations, and, separately, also examined the effects of reducing or increasing the level of facilitation on synaptic transmission properties (Fig. 1B).

### Model implementation

Realistic values of various parameters for the input stimulus were chosen in accordance with *in vivo* CA3 spike recordings from awake, freely moving rodents^35^, which have been used in previous STP studies^19,44,47^. This dataset comprises inhomogeneous spike trains spanning a broad range of discharge frequencies (∼5-60 Hz) and burst sizes (∼3-30 spikes per burst), with typically long intervals (∼several sec) separating individual discharges. We modeled two specific implementations of the temporal signal *ϕ*(t) shaping presynaptic spiking activity: one describing frequency modulation of sporadic place field firing (rate remapping), and another wherein it represents the variable duration (with fixed spiking frequency) of individual place field passes. Non-zero instances of *ϕ* were sampled randomly from an appropriate dynamic range accordingly. For the variable frequency case, spike rate for every pass was sampled from the 6-60 Hz range, and the step size was set to Δt = 0.5 s which also gives a consistent range of spike numbers per burst (3-30 on average). Alternately, variable-duration passes were modeled with a fixed in-field firing rate of 30 Hz (the average discharge frequency from experimental recordings) and individual passes spanning .1-1 s, again giving between 3 and 30 spikes per burst on average. Further, to simplify estimation of information rates in this case, the step size was fixed at Δt = 0.1 s, and every time a place field pass was reckoned to occur, that step was assigned a variable number of spikes, based on the duration (value of *ϕ*) corresponding to that instance. Thus, variable duration bursts were rescaled to a constant step Δt; this is a valid approximation when estimating average information rates over very long times, provided that burst durations ≪ 1/*r*_*s*_, which is compatible with the available data. To reduce errors in estimation of *R*_info_ in this approximation, facilitation and refilling time constants were also appropriately rescaled when implementing STP dynamics within every place field crossing by the corresponding duration *t*_*B*_ (*τ*_*f*_ → *τ*_*f*_ Δ*t*/*t*_*B*_ and *τ*_*r*_ → *τ*_*r*_Δ*t*/*t*_*B*_), to mitigate over (under) estimating the effect of STF (vesicle recycling).

Monte Carlo simulations of the STP model were carried out for a range of input rates (*r*_*s*_ = .05-.2 *s*^−1^), low noise levels (*r*_*n*_ = 0-1 *s*^−1^) and maximum RRP sizes (*N*_max_ = 1-15), and dependence of results on the basal 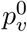 was characterized over ∼4 orders of magnitude 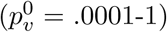. For every distinct parameter combination, 20 independent runs (each of 3 × 10^4^ s duration) were simulated, and the time-averaged rates of information flow and release events were estimated for every trial. The averages from our simulations were found to provide accurate estimates of the asymptotic information rates, justifying comparisons between different synaptic configurations in terms of the corresponding across-trial averages. To specifically assess the differential effect of short-term facilitation on synaptic function, every synapse with STF (referred to as a dynamic synapse) was compared with an equivalent static synapse which lacks facilitation while still exhibiting activity-dependent vesicle depletion (this corresponds to setting *α*_*f*_ = 0 in Eq. 3). In the following, only the results for the model with variable burst frequency are presented, although we have separately verified that the findings are closely reproduced for the variable-duration model as well.

All simulations, data analysis and visualization were performed in Python using the NumPy, SciPy and Matplotlib modules.

## 3. RESULTS

### 3.1 Improved signaling at unreliable synapses with short-term facilitation

How does short-term facilitation shape the vesicle code conveying information about presynaptic cell activity at stochastic hippocampal synapses? Experimental measurements reveal considerable diversity in the RRP size, release probability and STP properties across individual CA3-CA1 synapses^30,33^ (Fig. 1B). To explore the role of various synaptic attributes in modulating its transmission properties, we simulated the synaptic response to regular presynaptic spike trains occurring at different rates. Fig. 1C shows the response of a synapse to persistent spiking at 30 Hz (the average frequency in experimentally recorded bursts) for high and low initial failure rates (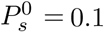 and 0.9) and a canonical maximum RRP size of *N*_max_ = 8. For the low *P*_*s*_ synapse, the time-dependent release probability initially increases due to strong activity-dependent facilitation and reaches a maximum, before the effect of vesicle depletion and slower recovery takes over, causing a drop in the response which eventually settles at a steady state value determined by the firing frequency. In contrast, the high *P*_*s*_ synapse shows a monotonically decreasing response with time, as it undergoes weaker facilitation and a larger initial *P*_*s*_ also implies faster depletion of the readily-releasable vesicle pool. The above qualitative differences illustrate the regimes of synaptic enhancement and depression encompassed by the STP model (Fig. 1A) that is based on physiological parameters for facilitation and depletion. To further bring out the differences between these two limits, we quantified the asymptotic/steady-state response amplitude of the dynamic synapse to input trains spanning a wide range of frequencies (0.1-100 Hz) as a function of the initial transmitter release probability. Fig. 1D shows the normalized synaptic response profiles for different base synaptic failure rates at a fixed RRP size (*N*_max_ = 8). High *P*_*s*_ synapses are most effective at transmitting spikes arriving at low frequencies, and with increasing facilitation (lower *P*_*s*_), the optimal transmission frequency is shifted to higher frequencies, demonstrating a transition from depression-dominated to facilitation-dominated behavior governed by the overall nature of STP dynamics (Eq. 3).

It is to be noted that RRP size also influences synaptic behavior along with the value of 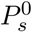, the latter being a function of the maximum number of vesicles available for release as well as the basal per-vesicle fusion probability. To demonstrate the role of RRP size in modulating synaptic behavior, the frequency-response relation estimated for different numbers of release-ready vesicles with a fixed basal synaptic failure rate 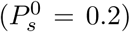 is shown in Fig. 1E. These curves highlight the role of RRP size in tuning the response profile of the synapse: synapses with a given fixed average rate of failures 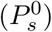 but differing in their number of available vesicles show a range of responses, from low-pass filtering for smaller *N*_max_ (corresponding to high basal 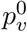) to higher optimal transmission frequencies for larger RRP sizes (lower basal 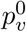). Taken together, the above examples (Figs. 1C-1E) capture the broad repertoire of behavior displayed across an ensemble of facilitating probabilistic synapses in the physiological regime controlled by the interplay among key synaptic parameters governing transmitter release and recovery.

Naturally occurring firing patterns in the CA3 region that encode behaviorally relevant integrated spatial and contextual signals are characterized by brief increases in firing frequency (spike bursts) separated by long periods of low activity^44^. We next examine synaptic processing of spike trains mimicking these activity patterns. Figure 2A illustrates the steps involved in our simulation of STP dynamics for a synapse with 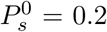 when the presynaptic spiking pattern carries information in the temporal sequence of burst occurrences *and* the variable firing frequency associated with every burst. The stochastic, time-varying signal is reflected in the brief spike discharges of the presynaptic neuron during passages through its preferred location (place field); this spiking activity drives the temporal dynamics of *p*_*v*_ and the transmitter release probability, eliciting a sequence of vesicle release events. We used a binning procedure to estimate the mean information content in the quantal release profile about the presynaptic signal (see Methods for details). Figure 2B (top left) shows the relative mutual information rate, *ℛ*_info_ (= *ℛ*_*rs*_/*ℛ*_*s*_), as a function of the basal probability of vesicle release 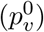 for different noise levels, for both a canonical synapse exhibiting STP (solid lines) and an equivalent synapse which does not show STF (dashed lines). Figure 2B (top right) illustrates the dependence of synaptic information rates on the RRP size for a fixed noise level of *r*_*n*_ = 0.1 Hz. These examples indicate a general enhancement of synaptic information transfer with short-term facilitation, as noted earlier in related contexts^19,48^. Further, this increase is more pronounced for synapses with lower release probability, aligning with our expectation that smaller basal 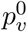 combined with stronger facilitation (Fig. 1B) accentuates the differential response to bursts and single spikes, enhancing the ability of the synapse to *selectively* transmit information-carrying spike discharges. To address the generality of the above results, we repeated our simulations over a biologically relevant range of parameter values (input rate = 0.05-0.2 Hz, noise level = 0-1 Hz and RRP size = 1-15). The overall difference in synaptic information capacity in the presence and absence of short-term facilitation is summarized as a distribution of relative changes in *ℛ*_info_ (percent difference of means) in Fig. 2B (bottom), and the color coding represents statistical significance of pairwise differences (two-sided Wilcoxon rank-sum test followed by Benjamini-Hochberg adjustment for multiple comparisons; blue: significant at FDR *<* .001, red: not significant). STF is found to robustly improve the fidelity of synaptic signaling in the physiological regime. The differential effect of STF scales inversely with the basal synaptic release efficacy, and is more marked for higher synaptic failure rates (lower 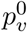). The effect of facilitation is diminished with increase in the basal release probability, and there is little difference in transmission efficacy between the dynamic and static synapses for 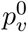 values above ∼0.1.

**FIG. 2:**
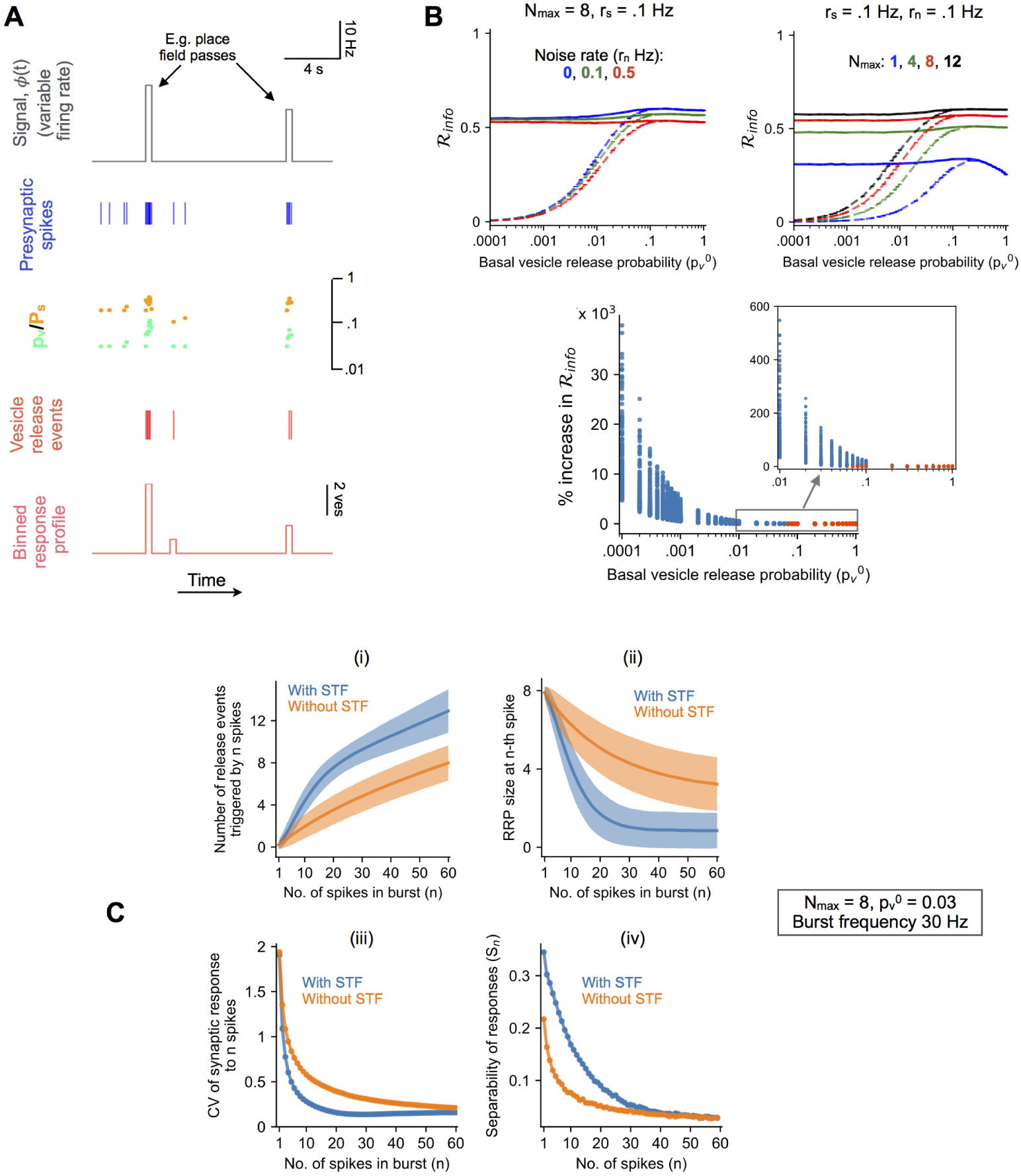
Elevated transmission of information-carrying spike patterns with synaptic short-term plasticity. **(A)** Time trace illustrating the transformation of an input signal (*ϕ*(t)) into a sequence of synaptic release events governed by STP. 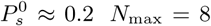, and *r*_*s*_ = *r*_*n*_ = 0.1 s^−1^. **(B)** *Top left* : Time-averaged rate of information transfer by synaptic release events (ℛ_info_) as a function of the basal per-vesicle fusion probability 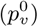 for a synapse with STF (continuous curves) and an equivalent synapse lacking STF (dashed curves). Different colors indicate different noise levels. (Results shown as mean *±* SEM over 20 independent simulations; *N*_max_ = 8 and *r*_*s*_ = .1 s.) *Top right* : Synaptic information transfer rates with STF (continuous curves) and without STF (dashed curves) for different numbers of available vesicles (different colors). (Results shown as mean *±* SEM over 20 independent trials; *r*_*n*_ = *r*_*s*_ = 0.1 s^−1^.) *Bottom*: Enhancement of synaptic information transmission with STF summarized as a distribution of relative changes (% difference of means relative to static synapse) over a biologically relevant range of input/model parameters (see Methods for details). Inset shows a magnified view of the .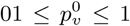 interval. Color-coding indicates statistical significance of pairwise differences (blue: significant at FDR *<* .001 level, red: not significant). **(C)** Synaptic response (i) and progressive depletion of the readily releasable vesicle pool (ii) as functions of the number of input spikes for a synapse with STF (blue) and lacking STF (orange). Response fluctuations are quantified in terms of the CV of number of release events (iii) and separability of responses to bursts differing in size by a single spike (iv). Results in (i) & (ii) shown as mean *±* SD (1000 independent trials); 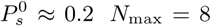, and spikes are Poisson-distributed with mean frequency of 30 Hz.

How can this improved reliability of synaptic information transmission be understood in simple terms? We recall that the synaptic response to information-carrying brief spike discharges consists of a variable number of release events; thus, the information content of the vesicle code in our formulation is essentially determined by how well the different output sizes (total number of releases triggered by a spike discharge) can discriminate between different input states (i.e. the variable spiking frequency, or duration, associated with every burst). We characterize the reliability of this mapping in terms of the cumulative number of released vesicles as a function of the burst size (number of spikes). Figure 2C (i) compares the responses of a canonical synapse (basal 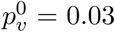) with and without STF to Poisson spiking at a mean rate of 30 Hz. In the presence of facilitation, not only is the average response amplitude (total number of release events for a given burst size) larger, but more importantly, it is also a more reliable readout of the burst size, due to reduced variability of responses relative to the static synapse. This difference is clearly seen in Figs. 2C (iii) & (iv), where two distinct measures for response fluctuations are plotted as functions of the number of spikes (*n*). The coefficient of variation (CV, defined as the standard deviation of the response relative to its mean) is lower for the synapse with STF, and the separability, defined as *S*_*n*_ = (*µ*_*n*+1_ - *µ*_*n*_)/(*σ*_*n*+1_ + *σ*_*n*_) (*µ*_*n*_ and *σ*_*n*_ denoting the mean and SD, respectively, of the response to *n* spikes), is larger with STF. The initial steeper increase in the mean response with the number of spikes (Fig. 2C (i)), together with smaller dispersion of responses (Figs. 2C (iii) & (iv)), implies better correspondence between the response amplitude and the burst length when STF is included. It is to be noted, though, that this increased reliability is also accompanied by faster depletion of the readily-releasable vesicle pool at the facilitating synapse (Fig. 2C (ii)), implying reduced dynamic range of burst sizes that can be conveyed by synaptic release events (this is indicated by a sharp change in the slope of the response profile for the STF synapse beyond some threshold spike number in Fig. 2C (i)). However, our results indicate that, in the biologically relevant parameter range considered here, STF, despite driving faster depletion, is distinctly advantageous for the shorter bursts of activity (Figs. 2C (iii) & (iv)), and leads to significant net improvement in synaptic information transfer (Fig. 2B). In sum, our simulations of STP dynamics highlight a crucial functional role for activity-dependent facilitation at unreliable hippocampal synapses, in enabling improved transmission of information represented by time-varying presynaptic cell activity in a physiologically relevant setting.

### 3.2 Reliable signaling at realistic STP synapses is nearly independent of their basal release properties

The results in the previous section (Fig. 2B) indicate that, for realistic number of vesicles, the signaling capacity of an STP synapse is not only increased relative to a static synapse, but, notably, also independent of its basal per-vesicle release rate 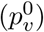 for the characteristic inhomogeneous spiking patterns associated with CA3 pyramidal cells studied here. To elaborate on the dependence of synaptic information transfer on its basal transmission probability, we scaled the mean information rate estimated for each value of 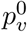 by the maximum value attained across the full range of 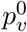 values considered (this was done separately for every combination of input rate, noise rate and vesicle pool size). This rescaling factors out the dependence on the other model parameters, and reveals the general trend in dependence of synaptic information transfer on its intrinsic reliability.

Figure 3A (top) shows the distribution of scaled information capacity values separately for synapses with STF (blue points) and lacking STF (grey points); each point represents a particular combination of model parameters and 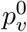. These results indicate that synaptic information transduction is robust to differences in the basal *p*_*v*_ at dynamic synapses. In other words, STP ensures that synapses with widely varying basal fusion probabilities transmit at comparable rates (median of values for each 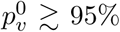 over the full range of 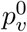 values considered here). In sharp contrast, information transfer at synapses lacking STF shows strong dependence on the magnitude of the basal *p*_*v*_, and is strongly impaired for synapses with 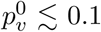.

**FIG. 3:**
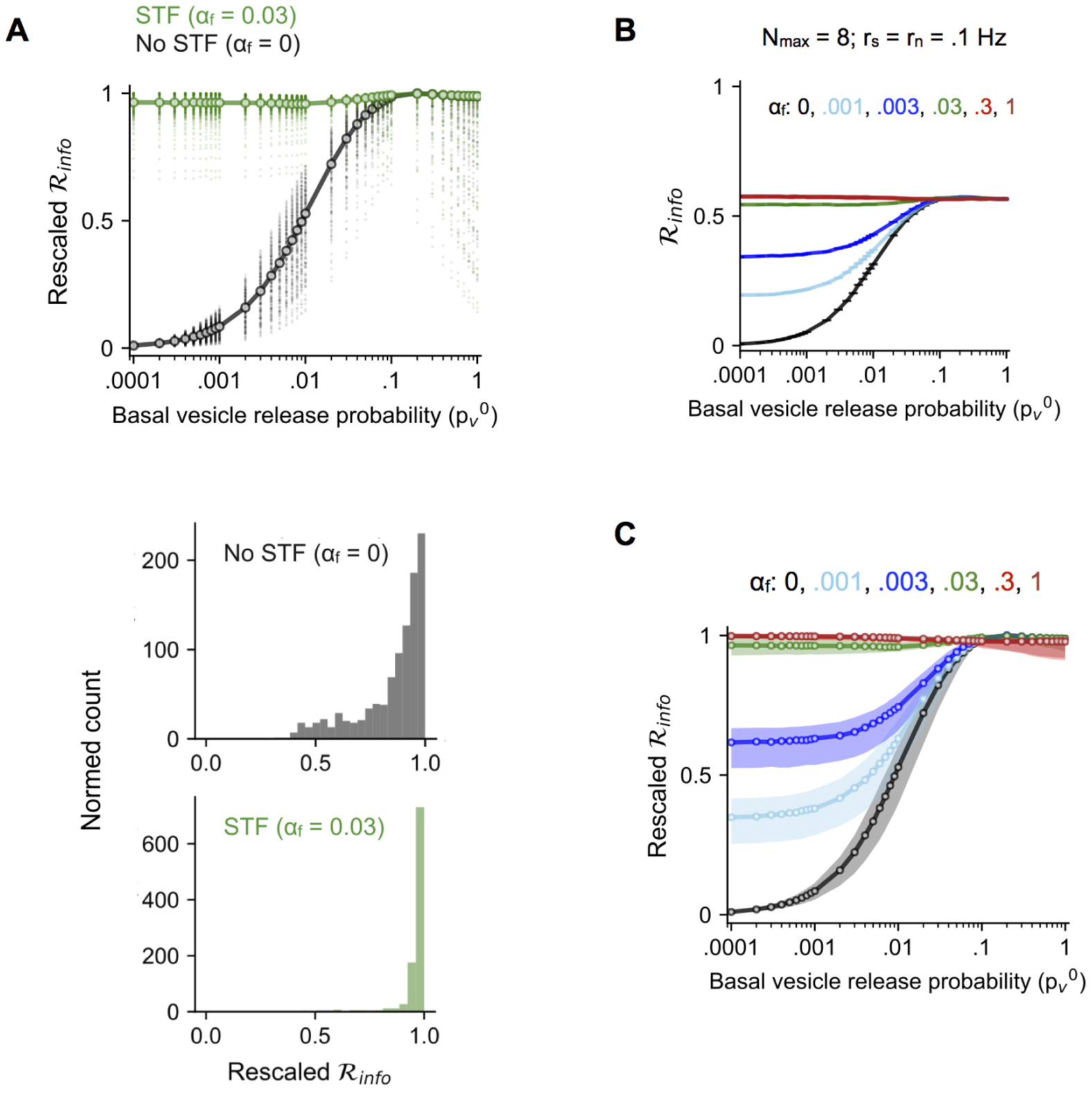
Information transmission properties in an ensemble of facilitating CA3-CA1 synapses. **(A)** *Top*: Dependence of information rate estimates (rescaled values) on the basal probability of vesicle release 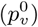 for synapses with STF in the biological regime (green) and lacking STF (black) over a realistic range of input/model parameters (see Methods for details). Every point represents a distinct parameter combination, and continuous lines connect the medians (one per value of *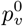*). *Bottom*: Distribution of rescaled information rates in a representative population of static (black) and facilitating (green) synapses with variable per-vesicle basal release probability 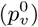 (n = 1000 synapses, randomly sampled from .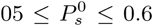 and 1 *≤N*_max_ *≤*15). **(B)** Estimated time-averaged information rate as a function of the basal per-vesicle release probability 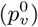 for synapses with different levels of facilitation (∼ 3 decades in the facilitation parameter *α*_*f*_). Results shown as mean *±* SEM (20 independent trials) for each choice of *α*_*f*_. *N*_max_ = 8; *r*_*s*_ = *r*_*n*_ = 0.1 s^−1^. The static synapse with no STF is shown in black. **(C)** Distributions of rescaled information rates over a realistic range of inputs/model parameters for different magnitudes of the synaptic gain *α*_*f*_ (0.001-1). Each distribution is displayed in terms of the medians and 25^*th*^-75^*th*^ percentile (interquartile) ranges (per value of 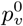).

These data are especially interesting in the light of the considerable heterogeneity in the magnitude of 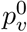 at hippocampal CA3 synaptic populations as reported by experimental studies^30–32^. Our analysis indicates that synaptic information transfer in the presence of dynamic gain control (Eq. 3) is nearly invariant to differences in the basal fusion probability per vesicle (Fig. 3A, bottom). We thus propose that physiologically realistic STP works to counteract degradation of presynaptic signals at synapses with small release probabilities, and enables these synapses to maintain stable information rates in the face of necessary heterogeneity in basal *p*_*v*_, arising from long-term changes associated with learning or homeostatic plasticity mechanisms on the circuit/network level.

The above analysis reveals significant overall difference in the nature of information transfer at stochastic synapses in the presence of short-term facilitation (Fig. 3A). How sensitive are these effects to its magnitude? Recalling that the dynamics of the release probability in our effective description of STP (Eq. 3) is essentially controlled by the gain parameter *α*_*f*_, which was adjusted to be compatible with experimental findings, we ask how the behavior of synapses changes for weaker or stronger facilitation. Fig. 3B shows an example of the (unscaled) synaptic information rate (mean *±* SEM) as a function of the basal 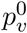 for a synapse with *N*_max_ = 8 vesicles, across a range of *α*_*f*_ values spanning ∼3 orders of magnitude (.001-1). Fig. 3C compares the distributions of rescaled information rate (estimated as before over a broad range of model parameters) for different levels of facilitation. The dispersion of estimates for each 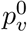 is represented in terms of the median (solid line) *±* interquartile range (IQR) separately for every *α*_*f*_. These results indicate that the profile of scaled synaptic information capacity is strongly modulated by changes in *α*_*f*_ especially at the smaller 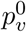 values (;≲ .1). This is a reflection of the greater sensitivity of facilitation at smaller basal release probabilities to changes in *α*_*f*_ in the STF model (Fig. 1B). In particular, reduction in *α*_*f*_ below the biological estimate 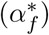 suppresses information transfer for smaller release probabilities and introduces heterogeneity in the ensemble behavior, whereas synaptic transmission capacity for 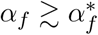 is nearly independent of the basal synaptic failure rate.

### 3.3 Short-term release dynamics regulates the capacity-cost trade-off at probabilistic synapses

In the previous section we have shown that short-term facilitation, in general, enables probabilistic synapses to signal the occurrence and length of brief high-frequency spike discharges more reliably. What is the theoretical limit on synaptic information capacity achievable at individual facilitating synapses, when transmitter release is governed by the STP model analyzed here (Fig. 1A)? For every combination of stimulus rate (*r*_*s*_), noise (*r*_*n*_) and RRP size (*N*_max_), we estimated the maximum rate of synaptic information transfer attainable when *α*_*f*_ and 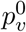 are allowed to vary, and we examined how well biological synapses (corresponding to 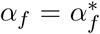) compare against this upper bound on *ℛ*_info_ (denoted as 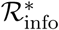). Fig. 4A displays the distribution of the normalized channel capacity for different choices of *α*_*f*_ (different colors); each point corresponds to a particular combination of model parameters and maximum RRP size. Our results show that biological synapses 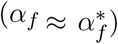 uniformly reach high, near-optimal, information rates under physiological conditions over ∼4 orders of magnitude of the basal per-vesicle release probability examined here (the median of normalized values for each 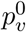 exceeds 90% over the full range of 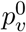 values considered). In contrast, probabilistic synapses with weaker facilitation, or no facilitation altogether, are much less effective at conveying information about presynaptic spiking activity, and the fidelity of information transfer at these synapses is markedly suppressed for 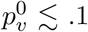 (Fig. 4A).

**FIG. 4:**
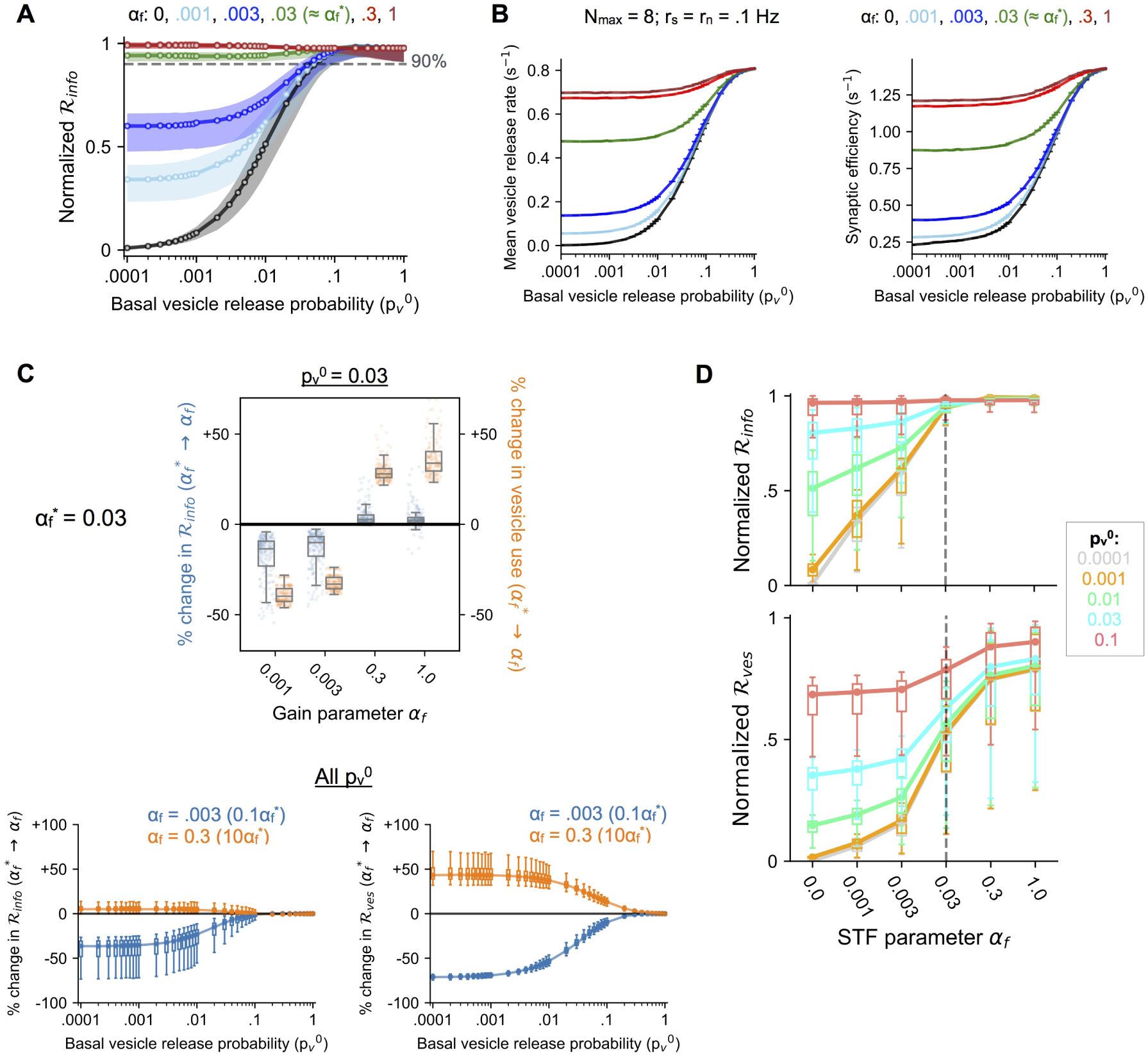
Optimal signaling and synaptic energy efficiency with physiologically realistic STP dynamics. **(A)** Distributions of normalized synaptic information rate (fraction of maximum capacity) over a biologically relevant range of parameters for different choices of the facilitation parameter *α*_*f*_ (different colors). Each distribution is displayed in terms of the data medians and interquartile ranges over a wide range of 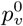 values. Realistic synapses 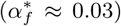 transmit at close to maximum capacity overall (median values *>* 90% across all 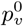). **(B)** Example profiles of the time-averaged rate of vesicle release (left) and synaptic energy efficiency (right) for synapses with different choices of *α*_*f*_ (all other parameter settings are same as in the example in Fig. 3B). Data shown as mean *±* SEM (20 independent simulations). **(C)** *Top*: Box-plots of relative changes (%) in synaptic information capacity and average vesicle requirement when synaptic gain is scaled either up or down by a factor of ∼ 10 relative to the physiological level 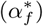 *for a synapse with basal 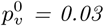*. Each distribution covers a biologically relevant range of input parameters and maximum RRP sizes (n = 180 points; see Methods for details). *Bottom*: Summary statistics of relative changes (%) in synaptic information rate (left) and vesicle usage (right) when *α*_*f*_ is scaled up (orange) or down (blue) by 10x, for a wide range of 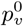 values. **(D)** Estimates of normalized signaling capacity (top) and vesicle use (bottom) as functions of the synaptic gain parameter *α*_*f*_ which spans ∼ 3 orders of magnitude (also shown, for reference, are results for the static synapse, corresponding to *α*_*f*_ = 0). Each box summarizes the results over a biologically relevant range of input/model parameters, and lines connect the median values for each choice of *α*_*f*_ ; profiles for different choices of the basal 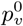 are represented by different colors. Vertical dashed lines highlight the biological set-point 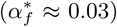.

Previous studies have emphasized the relevance of energetic constraints for a better understanding of neurobiological design on diverse scales^2,4^; examples from sensory systems, in particular, suggest that synaptic function may be significantly influenced by energy (resource) limitations^49,50^. To evaluate the potential role of energy constraints in shaping synaptic information processing in the hippocampus, we revisit the example in Fig. 3B, and quantify the synaptic vesicle use vis-à-vis information transfer at individual facilitating synapses. Fig. 4B shows the dependence of the average vesicle release rate and the energy efficiency of information transduction (∼ average number of vesicles needed to transmit a bit), respectively, on the basal *p*_*v*_ for different levels of synaptic facilitation (different colors) at a canonical CA3 synapse (*N*_max_ = 8). In general, vesicle use scales up with the basal probability of release 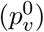 and with the strength of synaptic facilitation (*α*_*f*_), as expected (Fig. 4B, left). Notably, though, an increase in synaptic information transfer with stronger facilitation is accompanied by *reduction* in the synaptic energy efficiency, i.e., each released vesicle packs a smaller punch on average (Fig. 4B, right). The supralinear scaling of energy costs with synaptic information capacity implied by these examples suggests that in the context of realistic spiking patterns, individual CA3 synapses do not operate at optimal energy efficiency (according to the local measure of efficiency examined here), or minimize energy consumption; in fact, synapses lacking STP (black curves in Fig. 4B) require fewer releases per unit of information transmitted, albeit at significantly reduced overall information capacity, relative to dynamic synapses.

Do energy constraints, then, play no significant role in shaping the vesicle code at unreliable hippocampal synapses? Examining the regime of stronger facilitation 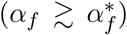 in the above example provides a potential clue in this regard. Figs. 3B and 4B together indicate that a canonical synapse operating in the physiological regime 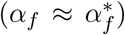 transmits information at near-optimal capacity, and further increase in *α*_*f*_ (by an order of magnitude, from 0.03 to 0.3 or 1) provides little additional benefit; the increased facilitation is, however, accompanied by a *disproportionately larger* increase in energy costs of synaptic transmission (this may be seen by comparing the green with the red/brown curves separately in Figs. 3B and 4B). This specific example suggests that biological CA3 synapses may be poised to operate near the upper bound on information transfer rate while energy usage is minimized to the extent that performance is not compromised.

To elaborate on the nature and generality of this energy-function trade-off, we compared biological STP synapses 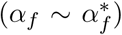 with synapses exhibiting weaker or stronger facilitation over ∼3 decades of magnitude, estimating the relative change in the mean signaling capacity and mean vesicle use per synapse when *α*_*f*_ is scaled up or down by a factor of ∼10 relative to its physiological reference value 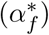. Fig. 4C (top) shows the distribution of relative changes over a range of model parameters (see Methods) for the specific example of 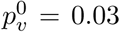, and each cluster of data points represents a different comparison 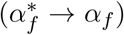. This aggregated data from our simulations indicates that stronger synaptic facilitation relative to the biological set-point provides little improvement in information transfer rates, but a *relatively larger* increase in energy use; reducing facilitation, on the other hand, is associated with a sharp reduction in synaptic information capacity. The overall differences evident in Fig. 4C (top) are found to be quite general and hold across a broad range of 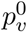 values examined (Fig. 4C, bottom).

The general trends suggested by Fig. 4C are displayed clearly in Fig. 4D, which shows how the normalized synaptic information capacity 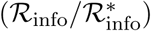 and normalized average rate of vesicle release 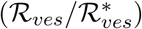 vary with the strength of synaptic facilitation (*α*_*f*_). Each box (median *±* IQR) summarizes the distribution of values for a particular 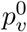, and the different colored lines connect the median values corresponding to each choice of 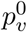. Simulations of our STP model suggest that synaptic information capacity is in general an increasing function of the strength of facilitation (*α*_*f*_), but saturates around the physiological level 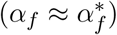 (Fig. 4D, top). Comparing it to Fig. 4D (bottom), further increase in synaptic gain comes at a larger energy cost, bringing diminishing returns. By contrast, reducing facilitation below the physiological operating point 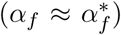 by ∼an order of magnitude compromises synaptic channel capacity considerably, and the suppression of information transfer rates is particularly marked at smaller basal vesicle release probabilities 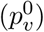. In sum, our results quantitatively demonstrate a novel form of local optimization embodied by short-term plasticity of vesicle release at probabilistic CA3-CA1 synapses, and suggest that, under physiological conditions, individual synapses do not consume more resources than necessary while supporting highest-possible fidelity of information transmission.

## 4. DISCUSSION

Is energy-efficient signaling a relevant design principle to account for salient properties of probabilistic transmitter release at individual hippocampal synapses? Previous investigations have focused on understanding energetic optimality at sensory pathway synapses^49–52^. Given the diversity in synaptic morphology and tight structure-function relationships in synapses observed across brain areas, questions on synaptic design must be specific and addressed in a local context. In line with this, we examined short-term plasticity at a cortical facilitating synapse, specifically the hippocampal Schaffer collateral-CA1 synapse. Our synaptic model invoked detailed characterizing properties of single CA3-CA1 terminals such as RRP size, release probability per vesicle and facilitation profiles derived from experiments and evaluated their impact on transmission of realistic activity patterns. This allowed us to obtain biologically relevant insights into synapse-specific design principles in the hippocampus. Our results provide a potentially normative account of biologically observed synaptic facilitation in terms of a local energy-information trade-off.

We estimated the capacity of a dynamic synapse, viewed as an unreliable channel, to communicate behaviorally relevant temporal signals coded in presynaptic spiking activity via discrete vesicle release events. Our quantitative analysis shows how short-term facilitation significantly improves the fidelity of synaptic information transduction. Remarkably, our simulations demonstrate that realistic STP enables CA3 synapses with vastly different basal release properties 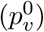 to convey brief, variable high-frequency spike bursts with comparable efficacy; in stark contrast, this invariance is absent at static or weakly facilitating synapses. Further, physiological information rates over a broad range of release probabilities closely approach the predicted maximum capacity of a facilitating synapse of this type, i.e. within the limits imposed by the overall form of the model of presynaptic dynamics analyzed here. We propose a nuanced form of optimality that is at odds with minimization of vesicle efficiency, quantified as the average number of quanta released per bit transmitted. Instead, our findings are consistent with the view that realistic STP synapses are poised, to within an order of magnitude in the gain parameter *α*_*f*_, to maintain near-maximal information transmission rates while penalizing excessive energy use (Fig. 5). Thus, we present evidence that energetic costs may also be important for regulating properties of activity-dependent facilitation at low-release probability synapses. Interestingly, an analogous form of optimality was previously proposed in the context of the mammalian visual system^53^. It was shown here that synaptic energy restrictions can significantly shape early stimulus representations in the retina, and that the observed centre-surround receptive fields provide the best balance between efficiency and performance, enabling near-maximal information transmission with largest possible synaptic energy savings. It remains to be seen, to what extent our findings are relevant to some of the other facilitating synapses in the mammalian central nervous system^54–56^.

**FIG. 5:**
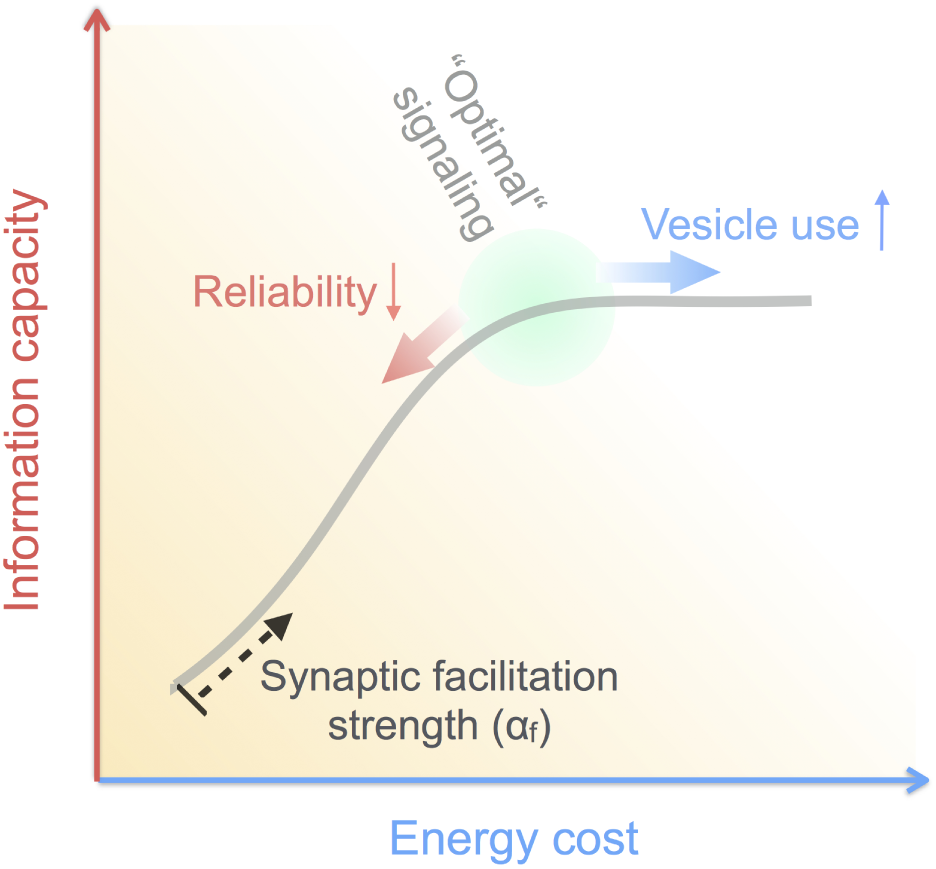
Activity-dependent short-term facilitation regulates the cost versus capacity trade-off at unreliable CA3-CA1 synapses. The feasible “configurational space” of an STP synapse (gray) is parametrized by the strength of synaptic gain, which constrains the relation between information transmitted across the synapse and the corresponding vesicle consumption. Our results suggest that biological synapses localize to the optimal regime indicated in green.

A key insight from our model is that synaptic information rates with physiological STF are nearly invariant to differences in the basal fusion probability per vesicle 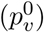 that are present among individual CA3-CA1 synapses. This synaptic diversity, on the one hand, may represent the intrinsic, across-synapse differences in the ultrastructural details regulating transmitter release^57^. On the other hand, a heterogeneous distribution of release probabilities may be a reflection of synaptically encoded memories, which are thought to be stored as distributed patterns of synaptic strength changes via activity-dependent long-term plasticity^58,59^. Experimental evidence, besides theoretical considerations, suggests that both Hebbian and heterosynaptic plasticity in the hippocampus can have a presynaptic as well as postsynaptic locus of expression^60–64^, that may be instantiated as persistent changes in 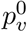. Additionally, variation in the basal *p*_*v*_ may arise as a consequence of homeostatic^65–67^ or neuromodulatory^68,69^ regulation of presynaptic calcium influx. In summary, several ongoing processes likely underlie the observed dispersion in CA3 presynaptic efficacies. Our analysis suggests that realistic STP dynamics operates at a set-point that compensates for this synaptic heterogeneity to support stable information rates, thereby implying a plausible mechanism to dissociate the dynamic synaptic interactions shaping ongoing circuit computations from the slower, longer-term adaptive changes that may be happening at these synapses due to learning or homeostatic adjustments.

Rate of vesicle release versus information transmitted is constrained by the synaptic gain parameter *α*_*f*_, which decides the operating point of the ensemble of CA3 synapses (Figs. 4D & 5). *α*_*f*_ may be tuned over evolutionary timescales to some suitable optimum determined by the relative influence of different, competing selective pressures. This aligns with recent understanding of the evolutionary diversification of the synaptic proteome that may have contributed to functional specializations in brain areas and emergent behavior^70^. In the context of the biophysical machinery governing transmitter release, what does the parameter *α*_*f*_ correspond to? Basal probability of spike-evoked release is governed by synchronous activation of the fast calcium sensor, Synaptotagmin-1 (Syt1)^71,72^. On the other hand, recent findings have identified a separate calcium sensor, Syt7, carrying a high-affinity binding site for Ca^2+^ but with relatively slower kinetics^73^, which was shown to be essential for progressive synaptic facilitation at CA3-CA1 terminals during persistent stimulation, but not for the initial (basal) synaptic response^74^. Efficacy of its interaction with the protein machinery mediating vesicle fusion, or the kinetic parameters governing its sensitivity to calcium, could thus provide a possible biophysical basis to interpret the parameter *α*_*f*_. Alternately, kinetic parameters regulating calcium-induced calcium release from intracellular stores which have been implicated in enabling short-term facilitation at hippocampal synapses^75,76^, or developmental parameters regulating the relative arrangement of calcium channels and the release machinery^77,78^, may account for the magnitude of *α*_*f*_. Biophysically detailed computational models of presynaptic calcium dynamics^79^, outside the scope of the present study, can potentially shed more light on the molecular underpinnings of STP approximated by the reduced description in Eq. 3 and help suggest physical interpretations of *α*_*f*_.

Although our study specifically examines the role of local constraints in shaping synaptic release properties, it is expected that synapse design also carries imprints of selection pressures at higher levels of neural organization. A number of previous studies have elaborated on the functional implications of synaptic short-term plasticity for collective dynamics on neuronal networks^80–82^. It is thus plausible, and quite likely, that properties of individual synapses reflect such system-level design considerations as well. The present work, in particular, does not account for the typical RRP size of CA3 synapses, which is experimentally found to be close to ∼10 vesicles per bouton^30,33^. Our analysis, in fact, indicates that average synaptic information capacity is a monotonically increasing function of the size of the RRP; thus, if information transfer is to be improved, larger synapses ought be favored, which runs somewhat counter to the limit on synapse size reported by experiments. We surmise that an optimal RRP size might represent a compromise between reliability of signaling at individual synapses and information processing capacity on the network level. Given strong constraints on neural volume (or equivalently, on total availability of synaptic resources) as proposed previously^4,83^, cortical connectivity might trade off high-fidelity synaptic transmission (proportional to RRP size) for increased network complexity from a higher density of smaller, albeit less reliable, synapses (based on scaling arguments)^84,85^. Detailed analysis of network information processing under physical constraints will be needed to evaluate the role of such an interaction across scales in shaping the design of fundamental computational elements in the brain.

To conclude, we propose that quantitative properties of probabilistic vesicle release at individual hippocampal synapses can be meaningfully interpreted in terms of a local cost-versus-capacity trade-off. Our results suggest that design of single synapses is primarily constrained to ensure optimal performance for diverse synaptic strengths, and this is achieved in an energetically cost-effective manner.

## Acknowledgments

This work was supported by IISER, Pune and received financial assistance from Science & Engineering Research Board, India (GM) and Wellcome Trust/DBT India Alliance (SN). GM acknowledges useful discussions on an earlier version of this work at the OIST Computational Neuroscience Course 2018.

## References

1 D. Attwell and S. B. Laughlin, An energy budget for signaling in the grey matter of the brain, Journal of Cerebral Blood Flow & Metabolism 21, 1133 (2001).

2 S. B. Laughlin, Energy as a constraint on the coding and processing of sensory information, Current opinion in neurobiology 11, 475 (2001).

3 V. Balasubramanian, D. Kimber, and M. J. B. Ii, Metabolically efficient information processing, Neural computation 13, 799 (2001).

4 S. B. Laughlin and T. J. Sejnowski, Communication in neuronal networks, Science 301, 1870 (2003).

5 A. Hasenstaub, S. Otte, E. Callaway, and T. J. Sejnowski, Metabolic cost as a unifying principle governing neuronal biophysics, Proceedings of the National Academy of Sciences 107, 12329 (2010).

6 J. G. G. Borst, The low synaptic release probability in vivo, Trends in neurosciences 33, 259 (2010).

7 G. Deco, E. T. Rolls, and R. Romo, Stochastic dynamics as a principle of brain function, Progress in neurobiology 88, 1 (2009).

8 W. B. Levy and R. A. Baxter, Energy-efficient neuronal computation via quantal synaptic failures, Journal of Neuroscience 22, 4746 (2002).

9 J. J. Harris, R. Jolivet, and D. Attwell, Synaptic energy use and supply, Neuron 75, 762 (2012).

10 R. S. Zucker and W. G. Regehr, Short-term synaptic plasticity, Annual review of physiology 64, 355 (2002).

11 M. V. Tsodyks and H. Markram, The neural code between neocortical pyramidal neurons depends on neurotransmitter release probability, Proceedings of the national academy of sciences 94, 719 (1997).

12 V. Matveev and X.-J. Wang, Differential short-term synaptic plasticity and transmission of complex spike trains: to depress or to facilitate? Cerebral Cortex 10, 1143 (2000).

13 J. Z. Tsien, P. T. Huerta, and S. Tonegawa, The essential role of hippocampal ca1 nmda receptor– dependent synaptic plasticity in spatial memory, Cell 87, 1327 (1996).

14 A. Gruart, M. D. Muñoz, and J. M. Delgado-García, Involvement of the ca3–ca1 synapse in the acquisition of associative learning in behaving mice, Journal of Neuroscience 26, 1077 (2006).

15 J. Basu and S. A. Siegelbaum, The corticohippocampal circuit, synaptic plasticity, and memory, Cold Spring Harbor perspectives in biology 7, a021733 (2015).

16 S. J. Middleton and T. J. McHugh, Silencing ca3 disrupts temporal coding in the ca1 ensemble, Nature neuroscience 19, 945 (2016).

17 C. Allen and C. F. Stevens, An evaluation of causes for unreliability of synaptic transmission, Proceedings of the National Academy of Sciences 91, 10380 (1994).

18 L. E. Dobrunz and C. F. Stevens, Response of hippocampal synapses to natural stimulation patterns, Neuron 22, 157 (1999).

19 Z. Rotman, P.-Y. Deng, and V. A. Klyachko, Short-term plasticity optimizes synaptic information transmission, Journal of Neuroscience 31, 14800 (2011).

20 A. Manwani and C. Koch, Detecting and estimating signals over noisy and unreliable synapses: information-theoretic analysis, Neural computation 13, 1 (2001).

21 M. S. Goldman, Enhancement of information transmission efficiency by synaptic failures, Neural computation 16, 1137 (2004).

22 R. Rosenbaum, J. Rubin, and B. Doiron, Short term synaptic depression imposes a frequency dependent filter on synaptic information transfer, PLoS computational biology 8, e1002557 (2012).

23 M. Salmasi, A. Loebel, S. Glasauer, and M. Stemmler, Short-term synaptic depression can increase the rate of information transfer at a release site, PLoS computational biology 15, e1006666 (2019).

24 C. Zhang and C. S. Peskin, Improved signaling as a result of randomness in synaptic vesicle release, Proceedings of the National Academy of Sciences 112, 14954 (2015).

25 J. O’Keefe and J. Dostrovsky, The hippocampus as a spatial map: preliminary evidence from unit activity in the freely-moving rat. Brain Res 34, 171 (1971).

26 J. E. Lisman, Bursts as a unit of neural information: making unreliable synapses reliable, Trends in neurosciences 20, 38 (1997).

27 L. Abbott and W. G. Regehr, Synaptic computation, Nature 431, 796 (2004).

28 A. M. Thomson, Facilitation, augmentation and potentiation at central synapses, Trends in neurosciences 23, 305 (2000).

29 J. S. Dittman, A. C. Kreitzer, and W. G. Regehr, Interplay between facilitation, depression, and residual calcium at three presynaptic terminals, Journal of Neuroscience 20, 1374 (2000).

30 L. E. Dobrunz and C. F. Stevens, Heterogeneity of release probability, facilitation, and depletion at central synapses, Neuron 18, 995 (1997).

31 V. N. Murthy, T. J. Sejnowski, and C. F. Stevens, Heterogeneous release properties of visualized individual hippocampal synapses, Neuron 18, 599 (1997).

32 N. Holderith, A. Lorincz, G. Katona, B. Rózsa, A. Kulik, M. Watanabe, and Z. Nusser, Release probability of hippocampal glutamatergic terminals scales with the size of the active zone, Nature neuroscience 15, 988 (2012).

33 T. Schikorski and C. F. Stevens, Quantitative ultrastructural analysis of hippocampal excitatory synapses, Journal of Neuroscience 17, 5858 (1997).

34 C. F. Stevens and Y. Wang, Facilitation and depression at single central synapses, Neuron 14, 795 (1995).

35 A. A. Fenton and R. U. Muller, Place cell discharge is extremely variable during individual passes of the rat through the firing field, Proceedings of the National Academy of Sciences 95, 3182 (1998).

36 K. Allen, J. N. P. Rawlins, D. M. Bannerman, and J. Csicsvari, Hippocampal place cells can encode multiple trial-dependent features through rate remapping, Journal of Neuroscience 32, 14752 (2012).

37 R. M. Grieves, E. R. Wood, and P. A. Dudchenko, Place cells on a maze encode routes rather than destinations, Elife 5, e15986 (2016).

38 Y. Aoki, H. Igata, Y. Ikegaya, and T. Sasaki, The integration of goal-directed signals onto spatial maps of hippocampal place cells, Cell reports 27, 1516 (2019).

39 A. Olypher, P. Lánskỳ, and A. Fenton, Properties of the extra-positional signal in hippocampal place cell discharge derived from the overdispersion in location-specific firing, Neuroscience 111, 553 (2002).

40 A. A. Fenton, W. W. Lytton, J. M. Barry, P.-P. Lenck-Santini, L. E. Zinyuk, Š. Kubík, J. Bureš, B. Poucet, R. U. Muller, and A. V. Olypher, Attention-like modulation of hippocampus place cell discharge, Journal of Neuroscience 30, 4613 (2010).

41 M. L. Shapiro, H. Tanila, and H. Eichenbaum, Cues that hippocampal place cells encode: dynamic and hierarchical representation of local and distal stimuli, Hippocampus 7, 624 (1997).

42 S. Leutgeb, J. K. Leutgeb, C. A. Barnes, E. I. Moser, B. L. McNaughton, and M.-B. Moser, Independent codes for spatial and episodic memory in hippocampal neuronal ensembles, Science 309, 619 (2005).

43 T. M. Cover and J. A. Thomas, Elements of information theory (John Wiley & Sons, 2012).

44 V. A. Klyachko and C. F. Stevens, Excitatory and feed-forward inhibitory hippocampal synapses work synergistically as an adaptive filter of natural spike trains, PLoS biology 4, e207 (2006).

45 M. H. Hennig, Theoretical models of synaptic short term plasticity, Frontiers in computational neuroscience 7, 45 (2013).

46 Y. Cai, J. P. Gavornik, L. N. Cooper, L. C. Yeung, and H. Z. Shouval, Effect of stochastic synaptic and dendritic dynamics on synaptic plasticity in visual cortex and hippocampus, Journal of neurophysiology 97, 375 (2007).

47 U. Kandaswamy, P.-Y. Deng, C. F. Stevens, and V. A. Klyachko, The role of presynaptic dynamics in processing of natural spike trains in hippocampal synapses, Journal of Neuroscience 30, 15904 (2010).

48 J.-P. Pfister, P. Dayan, and M. Lengyel, Synapses with short-term plasticity are optimal estimators of presynaptic membrane potentials, Nature neuroscience 13, 1271 (2010).

49 S. B. Laughlin, R. R. d. R. van Steveninck, and J. C. Anderson, The metabolic cost of neural information, Nature neuroscience 1, 36 (1998).

50 J. J. Harris, R. Jolivet, E. Engl, and D. Attwell, Energy-efficient information transfer by visual pathway synapses, Current Biology 25, 3151 (2015).

51 B. James, L. Darnet, J. Moya-Díaz, S.-H. Seibel, and L. Lagnado, An amplitude code transmits information at a visual synapse, Nature neuroscience 22, 1140 (2019).

52 J. J. Harris, E. Engl, D. Attwell, and R. B. Jolivet, Energy-efficient information transfer at thalamocortical synapses, PLoS computational biology 15, e1007226 (2019).

53 B. T. Vincent and R. J. Baddeley, Synaptic energy efficiency in retinal processing, Vision research 43, 1285 (2003).

54 A. Thomson, J. Deuchars, and D. West, Single axon excitatory postsynaptic potentials in neocortical interneurons exhibit pronounced paired pulse facilitation, Neuroscience 54, 347 (1993).

55 P. P. Atluri and W. G. Regehr, Determinants of the time course of facilitation at the granule cell to purkinje cell synapse, Journal of Neuroscience 16, 5661 (1996).

56 D. A. Henze, L. Wittner, and G. Buzsáki, Single granule cells reliably discharge targets in the hippocampal ca3 network in vivo, Nature neuroscience 5, 790 (2002).

57 H. L. Atwood and S. Karunanithi, Diversification of synaptic strength: presynaptic elements, Nature Reviews Neuroscience 3, 497 (2002).

58 B. L. McNaughton and R. G. Morris, Hippocampal synaptic enhancement and information storage within a distributed memory system, Trends in neurosciences 10, 408 (1987).

59 E. I. Moser, K. A. Krobert, M.-B. Moser, and R. G. Morris, Impaired spatial learning after saturation of long-term potentiation, Science 281, 2038 (1998).

60 C. F. Stevens and Y. Wang, Changes in reliability of synaptic function as a mechanism for plasticity, Nature 371, 704 (1994).

61 S. S. Zakharenko, L. Zablow, and S. A. Siegelbaum, Visualization of changes in presynaptic function during long-term synaptic plasticity, Nature neuroscience 4, 711 (2001).

62 R. Enoki, Y.-l. Hu, D. Hamilton, and A. Fine, Expression of long-term plasticity at individual synapses in hippocampus is graded, bidirectional, and mainly presynaptic: optical quantal analysis, Neuron 62, 242 (2009).

63 M. Letellier, Y. K. Park, T. E. Chater, P. H. Chipman, S. G. Gautam, T. Oshima-Takago, and Y. Goda, Astrocytes regulate heterogeneity of presynaptic strengths in hippocampal networks, Proceedings of the National Academy of Sciences 113, E2685 (2016).

64 R. P. Costa, Z. Padamsey, J. A. D’Amour, N. J. Emptage, R. C. Froemke, and T. P. Vogels, Synaptic transmission optimization predicts expression loci of long-term plasticity, Neuron 96, 177 (2017).

65 T. Branco, K. Staras, K. J. Darcy, and Y. Goda, Local dendritic activity sets release probability at hippocampal synapses, Neuron 59, 475 (2008).

66 C. Zhao, E. Dreosti, and L. Lagnado, Homeostatic synaptic plasticity through changes in presynaptic calcium influx, Journal of Neuroscience 31, 7492 (2011).

67 G. W. Davis and M. Müller, Homeostatic control of presynaptic neurotransmitter release, Annual review of physiology 77, 251 (2015).

68 B. P. Bean, Neurotransmitter inhibition of neuronal calcium currents by changes in channel voltage dependence, Nature 340, 153 (1989).

69 L.-G. Wu and P. Saggau, Presynaptic inhibition of elicited neurotransmitter release, Trends in neurosciences 20, 204 (1997).

70 R. D. Emes and S. G. Grant, Evolution of synapse complexity and diversity, Annual review of neuroscience 35, 111 (2012).

71 M. Geppert, Y. Goda, R. E. Hammer, C. Li, T. W. Rosahl, C. F. Stevens, and T. C. Südhof, Synaptotagmin i: a major ca2+ sensor for transmitter release at a central synapse, Cell 79, 717 (1994).

72 R. Fernandez-Chacon, A. Königstorfer, S. H. Gerber, J. García, M. F. Matos, C. F. Stevens, N. Brose, J. Rizo, C. Rosenmund, and T. C. Südhof, Synaptotagmin i functions as a calcium regulator of release probability, Nature 410, 41 (2001).

73 T. Bacaj, D. Wu, X. Yang, W. Morishita, P. Zhou, W. Xu, R. C. Malenka, and T. C. Südhof, Synaptotagmin-1 and synaptotagmin-7 trigger synchronous and asynchronous phases of neurotransmitter release, Neuron 80, 947 (2013).

74 S. L. Jackman, J. Turecek, J. E. Belinsky, and W. G. Regehr, The calcium sensor synaptotagmin 7 is required for synaptic facilitation, Nature 529, 88 (2016).

75 N. J. Emptage, C. A. Reid, and A. Fine, Calcium stores in hippocampal synaptic boutons mediate short-term plasticity, store-operated ca2+ entry, and spontaneous transmitter release, Neuron 29, 197 (2001).

76 N. Singh, T. Bartol, H. Levine, T. Sejnowski, and S. Nadkarni, Presynaptic endoplasmic reticulum contributes crucially to short-term plasticity in small hippocampal synapses, bioRxiv, 431866 (2018).

77 S. Nadkarni, T. M. Bartol, C. F. Stevens, T. J. Sejnowski, and H. Levine, Short-term plasticity constrains spatial organization of a hippocampal presynaptic terminal, Proceedings of the National Academy of Sciences 109, 14657 (2012).

78 N. P. Vyleta and P. Jonas, Loose coupling between ca2+ channels and release sensors at a plastic hippocampal synapse, Science 343, 665 (2014).

79 S. Nadkarni, T. M. Bartol, T. J. Sejnowski, and H. Levine, Modelling vesicular release at hippocampal synapses, PLoS computational biology 6, e1000983 (2010).

80 A. Levina, J. M. Herrmann, and T. Geisel, Dynamical synapses causing self-organized criticality in neural networks, Nature physics 3, 857 (2007).

81 G. Mongillo, O. Barak, and M. Tsodyks, Synaptic theory of working memory, Science 319, 1543 (2008).

82 J. F. Mejias and J. J. Torres, Maximum memory capacity on neural networks with short-term synaptic depression and facilitation, Neural computation 21, 851 (2009).

83 L. R. Varshney, P. J. Sjöström, and D. B. Chklovskii, Optimal information storage in noisy synapses under resource constraints, Neuron 52, 409 (2006).

84 C. M. Newman, Memory capacity in neural network models: Rigorous lower bounds, Neural Networks 1, 223 (1988).

85 D. B. Chklovskii, T. Schikorski, and C. F. Stevens, Wiring optimization in cortical circuits, Neuron 34, 341 (2002).

